# Monosynaptic Inputs to Excitatory and Inhibitory Neurons of the Intermediate and Deep Layers of the Superior Colliculus

**DOI:** 10.1101/824250

**Authors:** Ted K. Doykos, Jesse I. Gilmer, Abigail L. Person, Gidon Felsen

**Affiliations:** Neuroscience Graduate Program, University of Colorado School of Medicine, Aurora, CO 80045 USA; Department of Physiology & Biophysics, University of Colorado School of Medicine, Aurora, CO 80045 USA

**Keywords:** Monosynaptic, Excitatory, Inhibitory, Superior Colliculus, Neuroanatomy, Sensorimotor, Rabies

## Abstract

The intermediate and deep layers of the midbrain superior colliculus (SC) are a key locus for several critical functions, including spatial attention, multisensory integration and behavioral responses. While the SC is known to integrate input from a variety of brain regions, progress in understanding how these inputs contribute to SC-dependent functions has been hindered by the paucity of data on innervation patterns to specific types of SC neurons. Here, we use G-deleted rabies virus-mediated monosynaptic tracing to identify inputs to excitatory and inhibitory neurons of the intermediate and deep SC. We observed stronger and more numerous projections to excitatory than inhibitory SC neurons. However, a subpopulation of excitatory neurons thought to mediate behavioral output received weaker inputs, from far fewer brain regions, than the overall population of excitatory neurons. Additionally, extrinsic inputs tended to target rostral excitatory and inhibitory SC neurons more strongly than their caudal counterparts, and commissural SC neurons tended to project to similar rostrocaudal positions in the other SC. Our findings support the view that active intrinsic processes are critical to SC-dependent functions, and will enable the examination of how specific inputs contribute to these functions.

## 1. INTRODUCTION

The superior colliculus (SC) is a highly conserved midbrain structure critical for orienting behavior (Basso and May, 2017), as well as other associated functions such as spatial attention (Krauzlis et al., 2013) and multisensory integration (Stein and Stanford, 2008). The SC is organized into a superficial visual layer, which receives projections from the retina (Apter, 1945) and descending inputs from the neocortex (Kawamura et al., 1974), and intermediate and deep layers (SC_id_) that receive widespread input from several cortical and subcortical regions (Sparks and Hartwich-Young, 1989). The SC_id_ is organized into a topographic map of movement space, whereby small amplitude orienting movements are encoded rostrally and larger amplitude movements are represented caudally (Robinson, 1972; Wang et al., 2015). While much of our understanding of the role of the SC_id_ during behavior originated with work in primates making saccades to visual targets (Goldberg and Wurtz, 1972a; b; Wurtz and Goldberg, 1972; Lee et al., 1988), other work across a wider range of species points to a broader involvement of the SC_id_ (or the optic tectum (OT) the nonmammalian homologue of the SC) in other orienting behaviors (Sparks, 1999). For example, SC_id_/OT activity encodes orienting movements of the head in cats (Guillaume and Pélisson, 2001), monkeys (Freedman et al., 1996; Corneil et al., 2002; Walton et al., 2007), owls (du Lac and Knudsen, 1991), frogs (Meyer and Sperry, 1973), and bats (Valentine et al., 2002). SC_id_/OT neural activity also controls limb movements in cats (Courjon et al., 2004, 2015), monkeys (Werner et al., 1997; Philipp and Hoffmann, 2014), and mice (Steinmetz et al., 2018) as well as full body orienting movements in goldfish (Herrero et al., 1998) and rodents (Felsen and Mainen, 2008; Stubblefield et al., 2013). In addition to its role in orienting to targets across a wide range of evolutionarily diverse species, the SC is also critical for producing escape behavior away from aversive stimuli (Dean et al., 1986, 1989; Sahibzada et al., 1986).

Alongside our understanding of the SC_id_’s roles in behavior, a great deal is also known about which brain centers project to the SC_id_ (Edwards et al., 1979; Sparks and Hartwich-Young, 1989; Wolf et al., 2015). Several studies have employed anterograde and/or retrograde tracers demonstrating SC_id_ afferents originating from cerebral cortex (Garey et al., 1968; Edwards et al., 1979; Fries, 1984), thalamic areas (Edwards et al., 1974, 1979; Graybiel, 1974; Grofová et al., 1978), cerebellar nuclei (Batton et al., 1977; Kawamura et al., 1982), and several mesencephalic regions (Hopkins and Niessen, 1976; Grofová et al., 1978; Edwards et al., 1979). Potential roles for individual SC_id_ afferents range from transmitting behaviorally-relevant information about visual input (frontal eye field (FEF): Segraves and Goldberg, 1987; Sommer and Wurtz, 2000, 2001; Wurtz et al., 2001; lateral interparietal cortex: Paré and Wurtz, 2001; Wurtz et al., 2001; V1: Liang et al., 2015), recent experience (FEF: Sommer and Wurtz, 2001; M2: Duan et al., 2019), and target value (substantia nigra *pars reticulata* (SNr): Handel and Glimcher, 2000; Basso and Wurtz, 2002; Sato and Hikosaka, 2002; Bryden et al., 2011), to more active roles such as saccade initiation (FEF: Schiller et al., 1980; Hanes and Wurtz, 2001) and cessation (cerebellum: Goffart et al., 1998).

While these and other studies point to an integrative role for the SC in mediating behavior (Wolf et al., 2015), the SC_id_ itself contains a variety of cell types, and in order to fully elucidate its functional circuitry we need to better understand its cell-type-specific inputs (Oliveira and Yonehara, 2018; Masullo et al., 2019). As a first step, we focused on inputs to excitatory and inhibitory SC_id_ neurons (“eSCNs” and “iSCNs,” respectively). The SC_id_ is composed of ~70% glutamatergic cells and ~30% GABAergic cells (Mize, 1992) each with projection patterns within and between SC layers, to the contralateral SC, and out of the SC (Pettit et al., 1999; Isa and Hall, 2009; Sooksawate et al., 2011; Ghitani et al., 2014), suggesting that the interactions between eSCNs and iSCNs may play a key role in the SC_id_ computations underlying orienting behaviors. Thus, a critical piece of understanding SC_id_ function lies in discovering the specific projection patterns to eSCNs and iSCNs. Recent technological advances in mouse transgenics (Branda and Dymecki, 2004) and transsynaptic tracers (Wickersham et al., 2007; Wall et al., 2010; Luo et al., 2018) have allowed us to probe the organization of microcircuits with greater specificity. Thus, we leveraged Cre-lox recombination in conjunction with a transsynaptic retrograde rabies virus tracer strategy to label monosynaptic inputs to eSCNs and iSCNs, as well as to a subset of brainstem-projecting eSCNs neurons thought to drive orienting movements (Sooksawate et al., 2005, 2008). We found that projection patterns differed to these populations, suggesting cell-type-specific input integration, which has important implications for SC_id_ function.

## 2. MATERIALS AND METHODS

### Animals

All procedures followed the National Institutes of Health Guidelines and were approved by the Institutional Animal Care and Use Committee at the University of Colorado Anschutz Medical Campus. Animals were housed in an environmentally controlled room, kept on a 12-hour light/dark cycle and had ad libitum access to food and water. Adult mice of both sexes were used in these experiments (n = 7 males; n = 5 females). All mice were adult C57BL/6 (including Jackson labs homozygous Vglut2-Cre (RRID: IMSR_JAX:028863, Vong et al., 2011) and heterozygous Gad2-Cre (RRID: IMSR_JAX:010802, Taniguchi et al., 2011)) bred in house.

### Viral injections

AAV1-EF1.Flex.TVA.mCherry (UNC Vector Core; Watabe-Uchida et al., 2012) and AAV9.Flex.H2B.GFP.2A.oG (Salk Gene Transfer, Targeting and Therapeutics Core; Kim et al., 2016) were co-injected (100 nL of each; vortexed together after combining in equal proportions; Fig. 1A) unilaterally into the SC_id_ of Vglut2-Cre, Gad2-Cre, and wild-type mice. After a three week incubation period, a second injection of EnvA.SADΔG.eGFP virus was made at the same location (Salk Gene Transfer, Targeting and Therapeutics Core; Wickersham et al., 2007; Wall et al., 2010; Kim et al., 2016; Fig. 1A). Rabies virus injections (400 nL) were made at a 20° angle relative to vertical to avoid labeling cells in the superficial SC. Injection coordinates were varied along the rostrocaudal and dorsoventral axes and were made at −0.75 mm or −0.8 mm with respect to the midline. Mice were then sacrificed after an additional week and prepared for histological examination.

**Figure 1.**
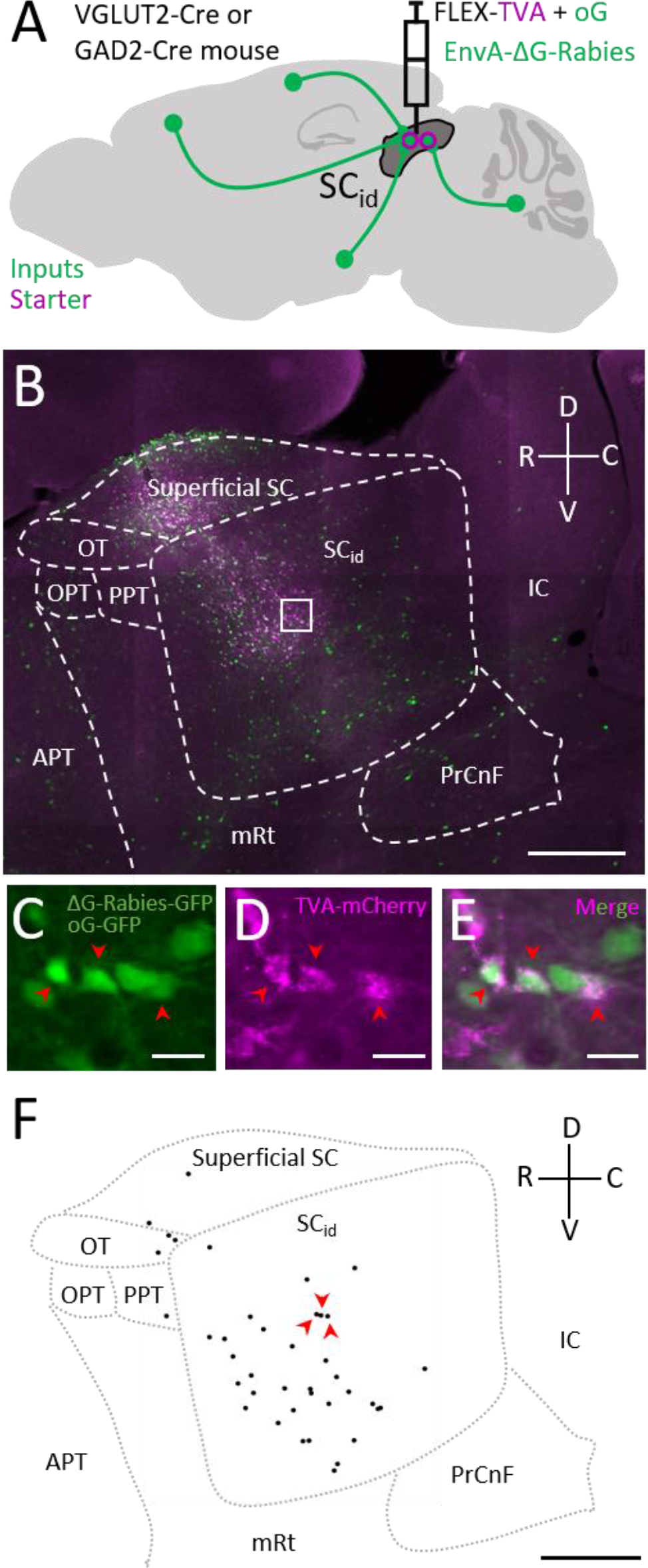
Rabies expression and identification of starter neurons. (A) Experimental strategy for targeting rabies virus to eSCNs and iSCNs. (B) Example image of an SC_id_ injection site. Green: oG-GFP and Rabies-GFP. Magenta: TVA-mCherry. White box indicates magnified area shown in C-E, and dashed outlines depict borders of the SC_id_ and surrounding brain areas. (C-E) Boxed area in B. Arrowheads indicate double-labeled starter neurons. (F) Dots indicate the location of the identified starter neurons displayed in panel B. Arrowheads indicate the starter neurons highlighted in C-E. Scale bars: 500 μm (B, F); 20 μm (C-E). Abbreviations: anterior pretectal nucleus (APT); inferior colliculus (IC); mesencephalic reticular formation (mRt); olivary pretectal nucleus (OPT); nucleus of the optic tract (OT); posterior pretectal nucleus (PPT); precuneiform area (PrCnF).

In experiments where rabies virus infection was targeted to SC_id_ output neurons, the above-described procedures were followed except wild-type mice were used and during the first surgery a retrograde Cre virus (CAV2-Cre; Peltékian et al., 2002) was injected into the medial pontine reticular formation (MPRF) contralateral to the injected SC_id_, a structure known to receive inputs from the SC_id_ (Huerta and Harting, 1982; Redgrave et al., 1990; Isa and Sasaki, 2002).

### Tissue preparation and imaging

Mice were overdosed with an intraperitoneal injection of a sodium pentobarbital solution, Pentobarbital (Sigma-Aldrich Inc.), and perfused transcardially with 0.9% saline followed by 4% paraformaldehyde. Brains were removed and postfixed for 4-24 hours then cryoprotected in 30% sucrose. Tissue was sliced in 40 μm serial sagittal sections using a freezing microtome and stored in 0.1 M phosphate buffered saline. Every third section was Nissl stained (Thermo Fisher Scientific, Cat# N21483, RRID: AB_2572212), mounted onto slides, and imaged in three colors using a slide-scanning microscope (Leica DM6000B Epifluorescence & Brightfield Slide Scanner; Leica HC PL APO 10× Objective with a 0.4 numerical aperture air; Objective Imaging Surveyor, V7.0.0.9 MT). Images were then converted to TIF files (OIViewer Application V9.0.2.0) for subsequent analysis. An additional set of images used for starter neuron analysis were acquired in sections near the injection site.

Images of input neurons displayed in Figure 2 were acquired with an Olympus IX81 with a disk scanning unit for confocal imaging (Olympus UPlanFL 20× Objective, 0.5 numerical aperture air) controlled with μManager software (https://micro-manager.org/, RRID: SCR_016865, Edelstein et al., 2010, 2014). Tile correction was performed in ImageJ (https://imagej.net/, RRID: SCR_003070, Rueden et al., 2017)/FIJI (http://fiji.sc, RRID: SCR_002285, Schindelin et al., 2012; Peng et al., 2017).

**Figure 2.**
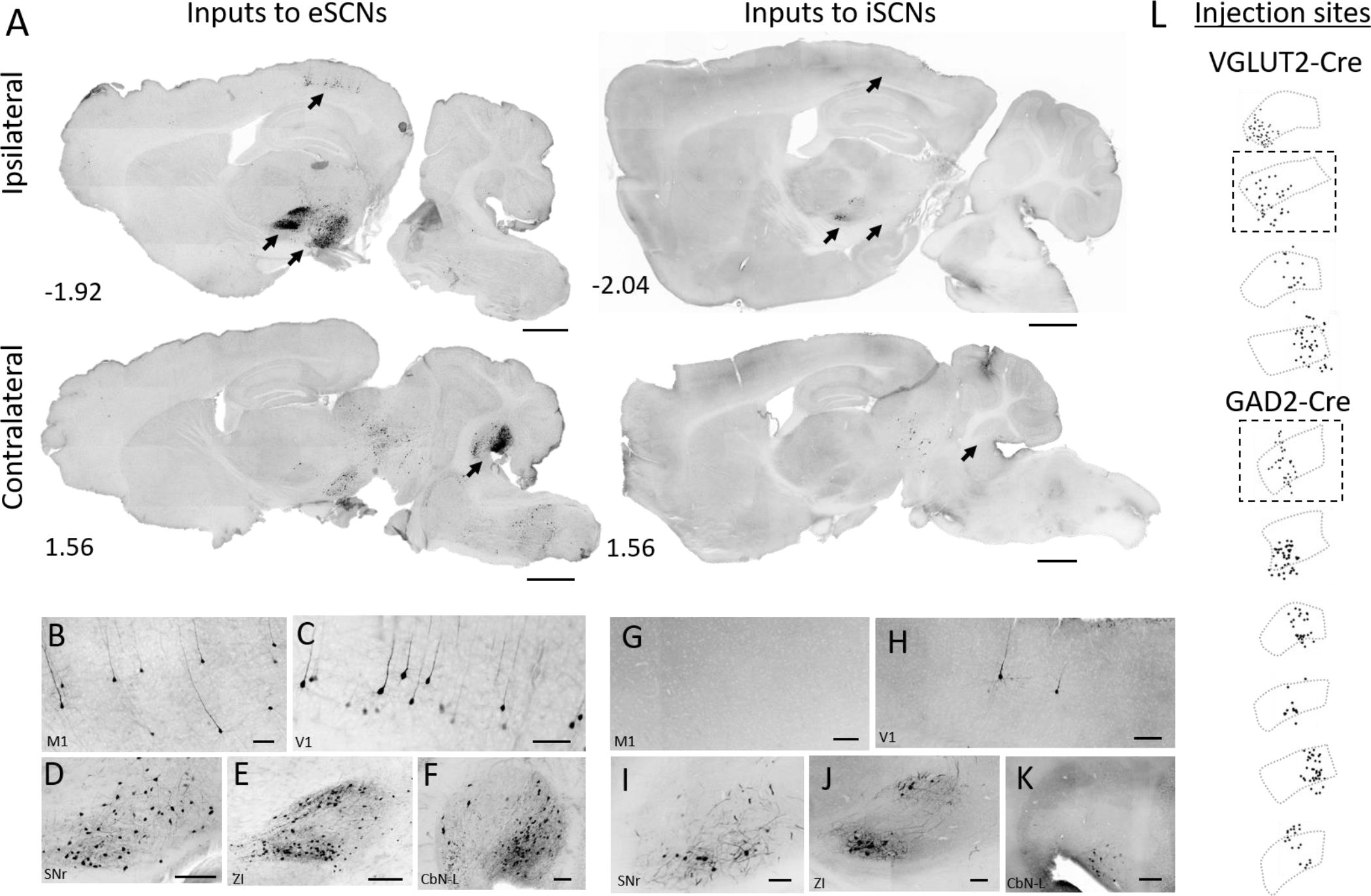
eSCNs receive more extrinsic inputs than iSCNs. Representative sagittal sections depicting inputs ipsilateral (top row) and contralateral (bottom row) to the injected SC_id_ in Vglut2-Cre (left column) and Gad2-Cre mice. Distance of section from midline (in mm) indicated by values in the lower left corner. (B-K) Representative images of input neurons to eSCNs (B-F) and iSCNs (G-K). Scale bars: 1 mm (A); 100 μm (B-K). (L) Injection sites depicting SC_id_ (gray dashed outline) and location of identified starter neurons (black dots) in a representative sagittal plane (between 0.36 mm and 1.08 mm lateral to midline) from each mouse. Dashed boxes indicate the two mice displayed in A-K.

### Starter neuron quantification and brain area classification

Rabies positive “starter neurons” were identified in ImageJ (FIJI) based on the following criteria: 1) the presence of GFP and mCherry, 2) visible neurites (Fig. 1C-E), and 3) fully overlapping mCherry-GFP signal. Neurons in which the mCherry signal extended beyond the borders of the GFP signal were not counted, since such labeling was more consistent with overlap of histone-tagged oG-GFP, not rabies-GFP, which readily fills cells.

Starter neuron coordinates were then exported to a MATLAB (http://www.mathworks.com/products/matlab/, RRID: SCR_001622) custom-written image viewer and classified as being within the SC_id_, the superficial SC, or any other region. Animals were only included in analyses if the majority of their starter neurons were located within the SC and the majority of those neurons were located within the SC_id_.

### Input neuron quantification and brain area classification

GFP-expressing input neurons were automatically identified using a semantic segmentation artificial neural network (SSN). The SSN was trained in MATLAB (Computer Vision System Toolbox) using custom-written scripts, to identify somata based on previously identified cell morphological data.

We used an initial training set of 4327 labelled images from 44 tissue sections obtained from 6 mice. 2581 of the sample images were images of known cells, and the remaining 1746 were images that were verified not to contain any cells, to provide examples of negative data. Each image was a 200 × 200 pixel image either centered on the cell coordinates, or arbitrarily chosen from the images of the sections. Tissue containing labeled and unlabeled neurons was used for semantic labeling in the training set. Cell body morphology was estimated by taking pre-identified soma coordinates and using the MATLAB ‘regionprops’ function to isolate features in the selected images. This process was optimized to find the human-identified cells within an image. If the cell body boundaries could not be resolved algorithmically, a circle with a radius of six pixels was drawn centered on the cell coordinates instead.

The extracted images were prepared for SSN usage using a custom MATLAB script, and the network was trained iteratively to optimize accuracy, using the following method: after each round of training and scoring, misses were saved as an image and replicated in the training data pool, and false positives were added to the negative image training pool. After several iterations, the final training dataset contained 16,443 images.

To validate that the network performed at an accuracy similar to a human observer, three experimenters who had not previously labelled images (A.P., G.F., and J.G.) performed the image coordinate identification process on 5 sample images. The agreement in labeling between the SSN and experimenters was comparable to inter-experimenter agreement (agreement between experimenters on detected cells: 69.1% ± 6.3% of all cells [standard deviation]; experimenter agreement with deep learning algorithm: 70.7% ± 8.3% [standard deviation]).

A quality control step was added to ensure that machine identified neurons were in agreement with human assessment. Each machine identified neuron was output as an image, and false positives were manually deleted. Thus, the final dataset included only neurons remaining after manual curation. The coordinates of SSN identified neurons were then exported to a MATLAB custom-written image viewer and classified according to brain area based on a standard mouse atlas (Paxinos and Franklin, 2013). We mainly focused on descending projections that were sufficiently far from the injection site to exclude contamination by starter neurons (See Results).

### Starter and input neuron analyses

Input neuron counts in each brain area were normalized to the number of SC_id_ starter neurons in that mouse to yield a measure of “projection strength”, a standard metric to correct for variability in viral expression (Watabe-Uchida et al., 2012; Sun et al., 2014). Subsequent analyses of projection strength were performed in MATLAB. Brain areas included in analyses comparing eSCNs to iSCNs had projection strengths to eSCNs or iSCNs > 0.01 (> 1 input neuron per 100 starter neurons). Areas with projection strengths to iSCNs > 0.005 were used in Figure 5C and 5D. The laterality preference in Figure 4 was computed as: LP = (2 × C) − 1, where C is the fraction of contralateral inputs. This yielded normalized values between −1 (strongest ipsilateral preference) and 1 (strongest contralateral preference). Random inputs in Figure 7F were obtained by averaging iterations of randomly generated input neuron positions (1000 iterations; 1707 random input neuron positions [eSCNs] or 145 random input neuron positions [iSCNs] per iteration).

The normalized rostrocaudal positions of starter and input neurons used in Figures 5 and 7 were obtained by setting the rostral-most and caudal-most extents of the SC_id_ in each section to 0 and 1, respectively, and determining the fractional location of each neuron along the rostrocaudal axis.

## 3. RESULTS

### Rabies expression and identification of starter neurons

We used a modified rabies virus, EnvA-ΔG-Rabies-GFP, to transsynaptically label neurons projecting monosynaptically to eSCNs in Vglut2-Cre (RRID: IMSR_JAX:028863, Vong et al., 2011) or iSCNs in Gad2-Cre mice (RRID: IMSR_JAX:010802, Taniguchi et al., 2011; Fig. 1A). Cell-type specificity of viral infection and spread was achieved using a Cre-dependent trans-complementation strategy (Wickersham et al., 2007; Wall et al., 2010; Watabe-Uchida et al., 2012; Kim et al., 2016; Beitzel et al., 2017). The majority of rabies “starter” neurons were located within the SC_id_ (Fig. 2B, F). Only cells with visible neurites that were both GFP+ and mCherry+ were classified as starter neurons (see Materials and Methods; Fig. 1C-E). Subsequent analyses were then used to identify input neurons and map them according to brain area (see Materials and Methods). Neuron counts in each area were then normalized to the number of SC_id_ starter neurons to yield a measure of “projection strength”, a standard metric to correct for variability in viral expression (Watabe-Uchida et al., 2012; Sun et al., 2014). Using this approach we quantified projection patterns to eSCNs and iSCNs.

### Extrinsic inputs to eSCNs and iSCNs

We found that eSCNs receive a greater number and a more diverse set of extrinsic inputs than iSCNs, even after accounting for the fact that eSCNs are more numerous than iSCNs. Strikingly, we observed much stronger projections to eSCNs than iSCNs, both across the brain (eSCN projection strength: 15.46 ± 6.56 [median ± median absolute deviation], n = 4 mice; iSCN projection strength: 0.73 ± 0.59 [median ± median absolute deviation], n = 6 mice; Wilcoxon rank sum test [one-tailed], p < 0.005; Fig. 2A; Fig. 3B), and within individual brain areas (Fig. 2B-K; Fig. 3A-B). With respect to individual brain areas, 36.73% (18/49) were found to send significantly stronger projections to eSCNs than iSCNs (Wilcoxon rank sum test [one-tailed]; p < 0.05; Fig. 3A). We also found that eSCNs tend to receive more inputs than iSCNs when input areas are grouped We also found that eSCNs tend to receive more inputs than iSCNs when input areas are grouped by developmentally-defined categories of telencephalon, diencephalon, mesencephalon, and cerebellum (Wilcoxon rank sum test [one-tailed]; p < 0.01; n = 4 mice [eSCNs]; n = 6 mice [iSCNs]; Fig. 3C).

**Figure 3.**
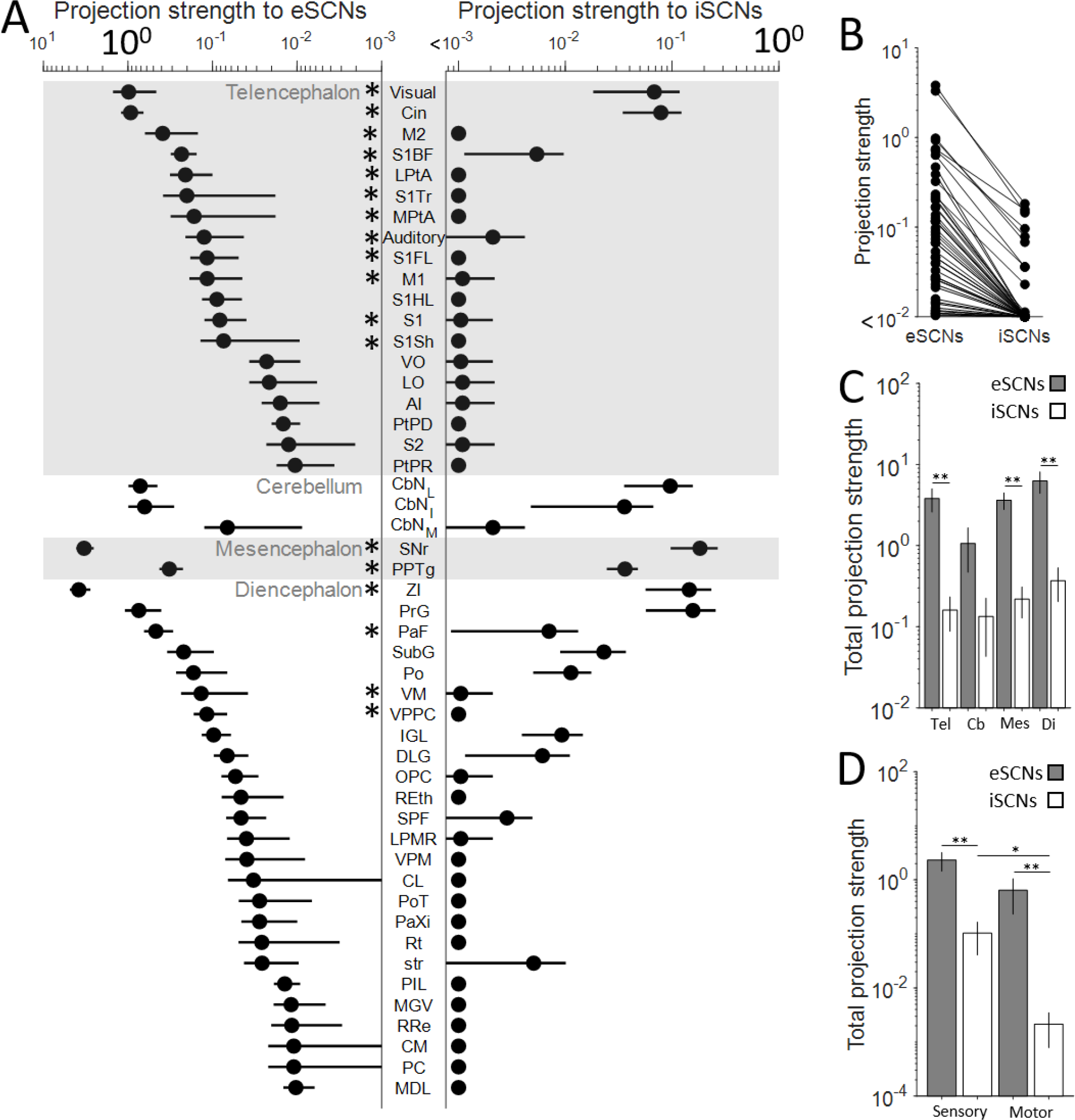
Projection strength to eSCNs and iSCNs. (A) Projection strength to eSCNs (left) and iSCNs (right) for each brain area. Note eSCNs and iSCNs are plotted on different scales for visibility. (B) Direct comparison between projection strength to eSCNs and iSCNs, shown on the same scale. (C) Total projection strength to eSCNs and iSCNs from developmentally defined brain regions. Tel (telencephalon); Cb (cerebellum); Mes (mesencephalon); Di (Diencephalon). (D) Total projection strength to eSCNs and iSCNs from regions of cortex and thalamus grouped according to sensory or motor function. Mean ± SEM (eSCNs: n = 4 mice; iSCNs: n = 6 mice). *: p<0.05; **: p<0.01.

eSCNs and iSCNs tended to receive their strongest projections from the same brain areas. The areas with the most prominent projections to the SC_id_ were zona incerta and SNr, although inputs from the pregeniculate nucleus of the prethalamus made up a substantial proportion (12%) of all inputs to iSCNs (Fig. 3A). Additionally, eSCNs and iSCNs both received their strongest cortical projections from visual and cingulate cortex and their strongest cerebellar projections from lateral and intermediate cerebellar nuclei (Fig. 3A).

Interestingly, when we classified cortical or thalamic brain areas as “sensory” or “motor” (Watson et al., 2012) we found that iSCNs were targeted by sensory areas more than motor areas (Wilcoxon signed rank test [two-tailed]; n = 6 mice; p = 0.0313; Fig. 3D); however, this was not true of eSCNs (Wilcoxon signed rank test [two-tailed]; n = 4 mice; p = 0.125; Fig. 3D). Taken together, these results suggest that eSCNs and iSCNs receive their strongest inputs from a similar set of brain areas, but that eSCNs receive a greater number and more diverse set of inputs overall.

### Laterality of inputs to eSCNs and iSCNs

Consistent with previous findings (Sparks and Hartwich-Young, 1989), the SC_id_ exhibited much stronger ipsilateral than contralateral input from most brain regions (Fig. 4; see Materials and Methods). Cerebellotectal projections deviated from this pattern, which is consistent with the well-established robust interconnectivity of the cerebellum with the contralateral side of the brain. The ipsilateral preferences from other areas were most strongly observed in inputs to iSCNs; however, input preference to eSCNs, specifically from cortical and thalamic areas, ranged from strongly ipsilateral to weakly contralateral (Fig. 4). For example, inputs to eSCNs from primary somatosensory cortex were almost entirely ipsilateral, while eSCN inputs from agranular insular cortex had a mild contralateral preference. Our observations indicate that overall, eSCNs receive more bilateral input than iSCNs, suggesting that eSCNs may be more involved in integrating sensory inputs from both sides of the body, while iSCNs may process more exclusively ipsilateral input.

**Figure 4.**
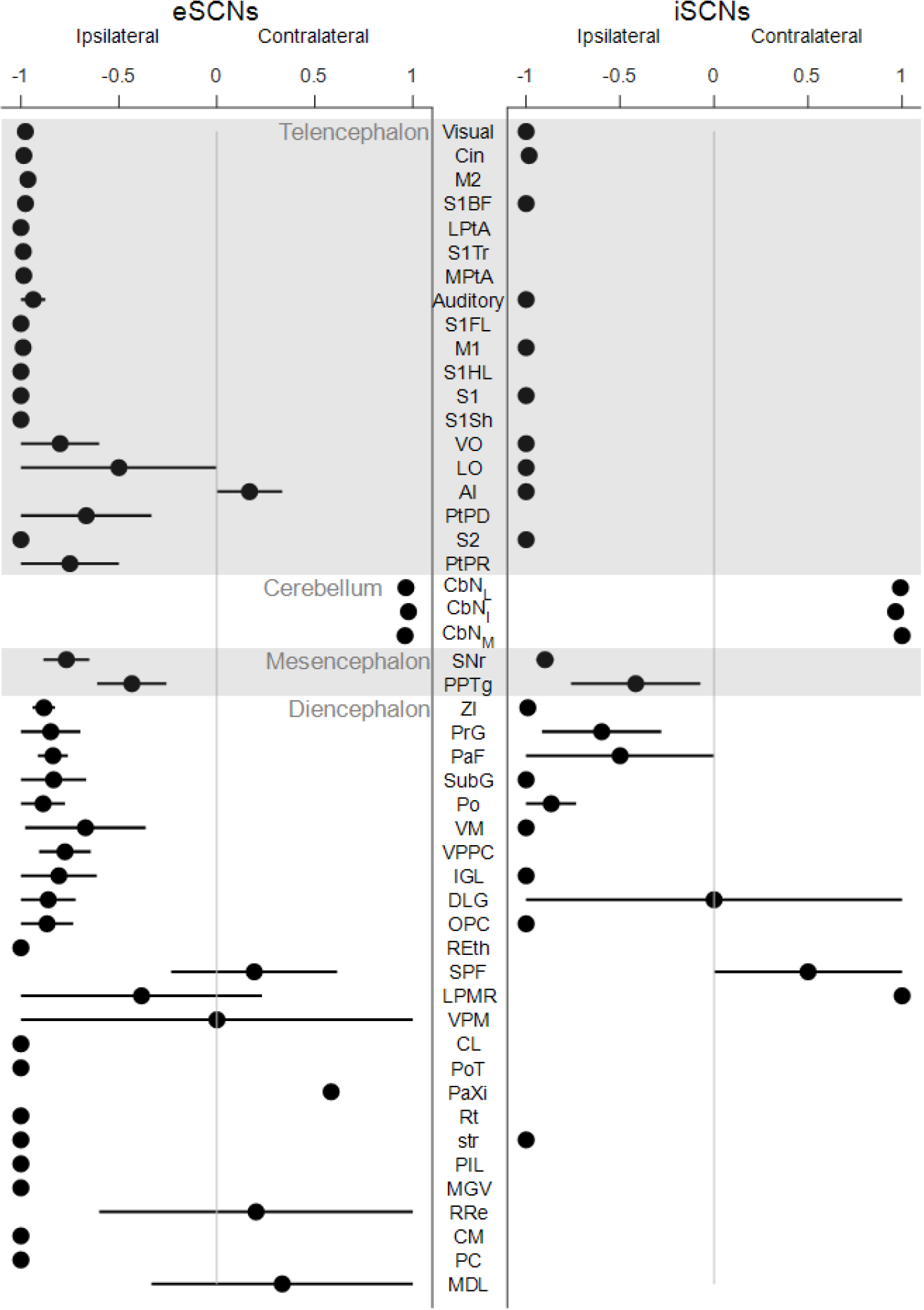
Laterality of SC inputs. Laterality preference to eSCNs (left) and iSCNs (right) for each brain area. Areas ordered as in Figure 3. Mean ± SEM (eSCNs: n = 4 mice; iSCNs: n = 6 mice).

### Relationship between projection strength and rostrocaudal position of starter neurons

The SC_id_ has a well-characterized topographic organization along its rostrocaudal axis, with small orienting movements encoded rostrally, and larger orienting movements represented caudally (Sahibzada et al., 1986; Gandhi and Katnani, 2011). We therefore examined the extent to which input neuron pattern depended upon the rostrocaudal position of starter neurons within the SC_id_. Across mice, we labeled both eSCNs and iSCNs at several points along the rostrocaudal axis (Fig. 5A). We observed a tendency across areas for stronger projection strengths to be associated with more rostrally located starter neurons: (eSCN: median Pearson Correlation Coefficient r = −0.36, p = 0.0024, n = 49 brain areas [Wilcoxon signed rank test]; iSCN median Pearson Correlation Coefficient r = −0.41, p = 0.0044, n = 16 brain areas [Wilcoxon signed rank test]; Fig. 5C, D). This trend was observed in both cell types. Overall, these findings indicate that eSCNs and iSCNs toward the rostral pole of the SC_id_ receive moderately more inputs than their caudal counterparts.

**Figure 5.**
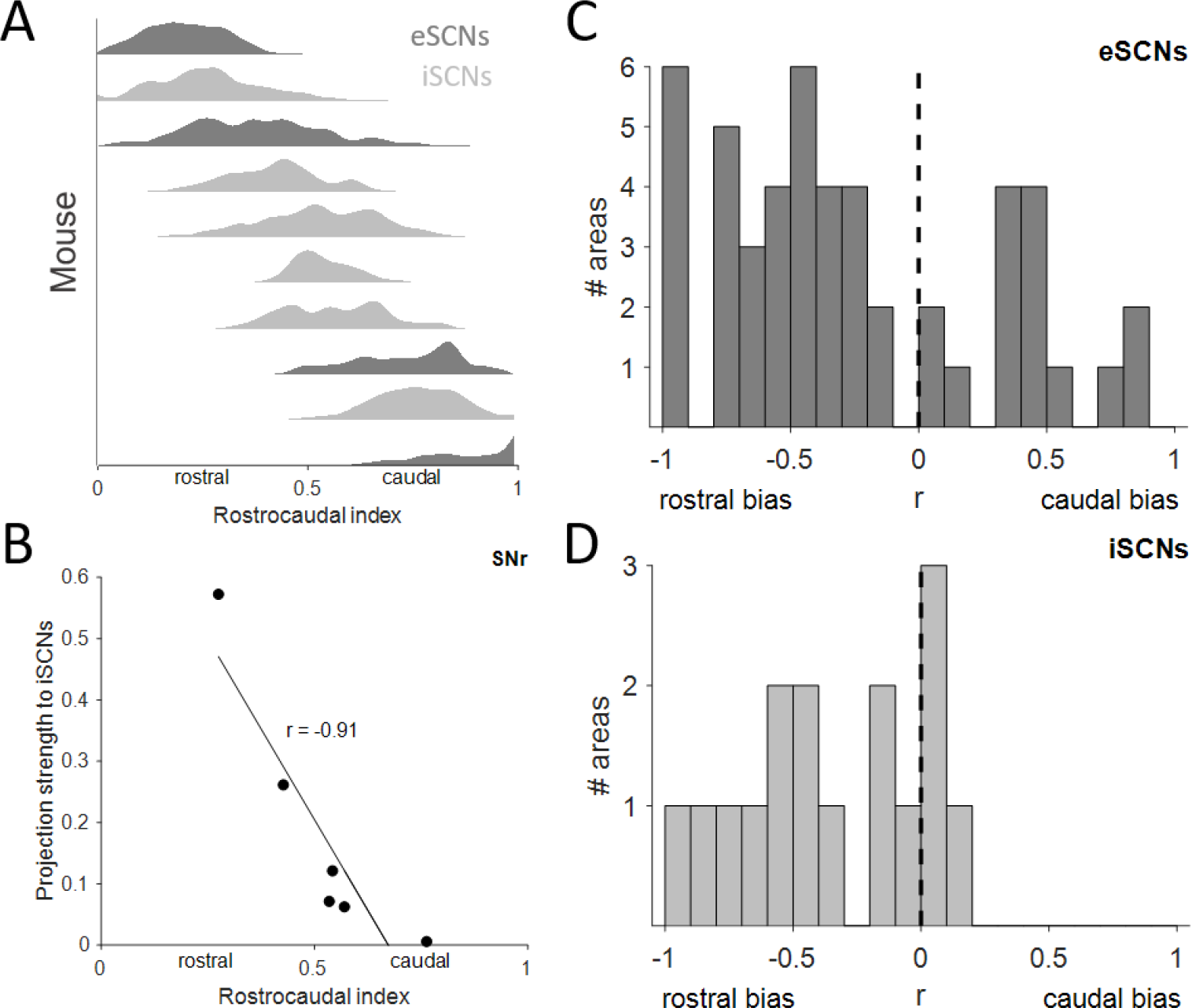
Extrinsic inputs favor the rostral SC. (A) Normalized rostrocaudal position of the starter neuron population for each mouse. (B) Pearson’s correlation between mean rostrocaudal position of starter neurons and projection strength from SNr to iSCNs. (C, D) Correlation coefficients relating mean projection strength to rostrocaudal index of injection, as in B, for all areas: eSCNs (C); iSCNs (D). Dashed line at x = 0 indicates no rostrocaudal bias.

### Extrinsic inputs to subset of tectofugal eSCNs

eSCNs comprise about 70% of SC_id_ neurons (Mize, 1992) and are diverse with respect to morphology and projection patterns (Pettit et al., 1999; Isa and Hall, 2009; Sooksawate et al., 2011; Ghitani et al., 2014). As a first step toward identifying patterns of inputs to putative subclasses of eSCNs, we focused on inputs to crossed tecto-reticular neurons (“CTRNs”) which are thought to be critical drivers of orienting movements and have been characterized in slice experiments (Sooksawate et al., 2005, 2008). We combined our Cre-dependent rabies trans-complementation approach with a retrograde Cre virus (Peltékian et al., 2002) injection into the contralateral MPRF of wild-type mice (see Materials and Methods; Fig 6A), such that starter neurons would be limited to MPRF-projecting SC_id_ neurons. We found that secondary motor cortex, the lateral cerebellar nuclei, SNr, and zona incerta were the only areas that provided measurable input to CTRNs (Fig. 6B), and that eSCNs received much stronger projections than CTRNs from these and other areas (Wilcoxon signed rank test [one-tailed]; p < 2 × 10^9^; n = 49 brain areas; Fig. 6C). There was no difference in the laterality preference to eSCNs and CTRNs. Together, these results indicate that, although CTRNs play a direct role in the orienting motor output function of the SC_id_, they receive a smaller and less diverse set of extrinsic inputs than the general eSCN population.

**Figure 6:**
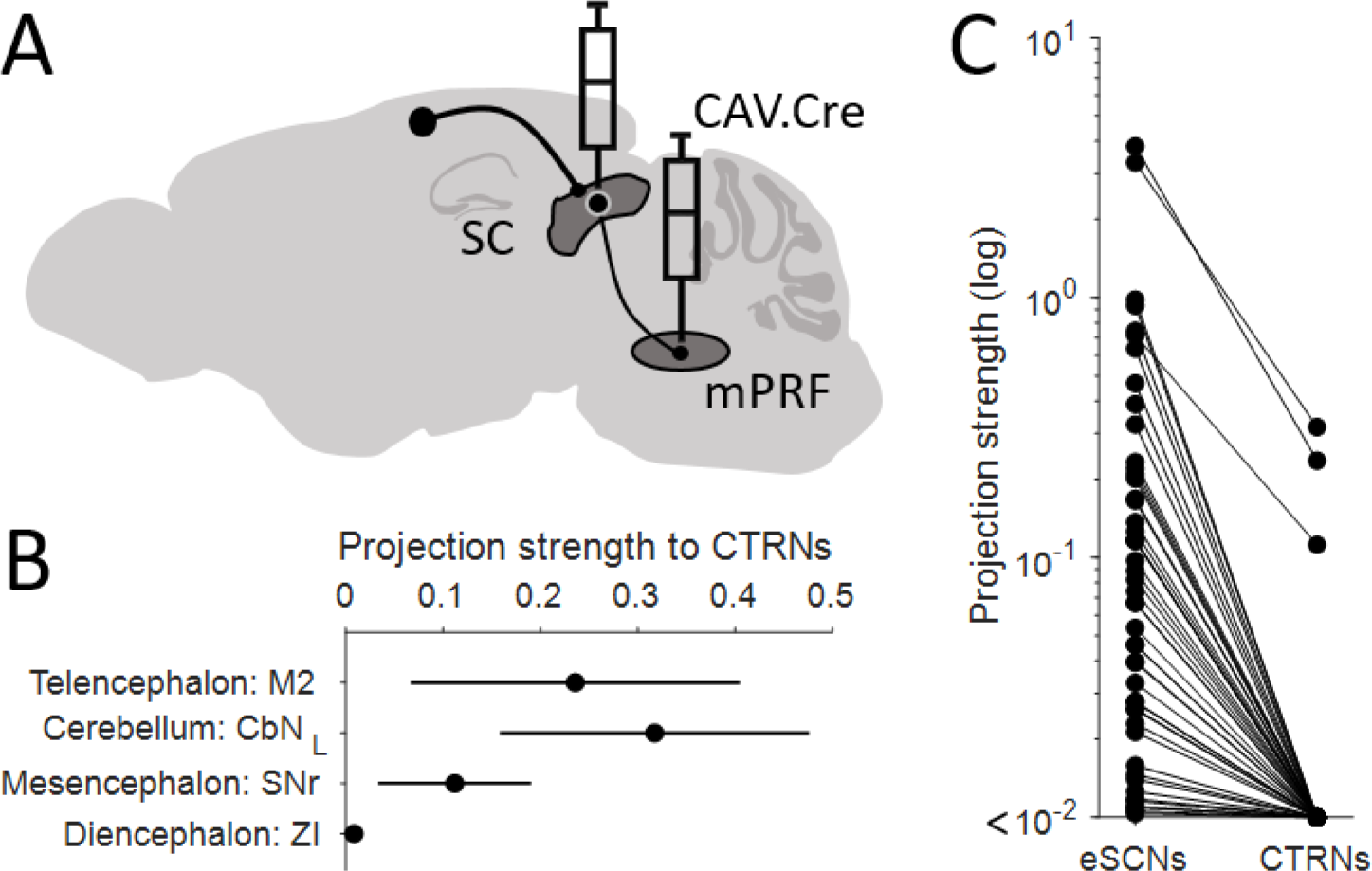
CTRNs receive fewer inputs than eSCNs. Experimental strategy targeting CTRNs. (B) Projection strength from areas targeting CTRNs. Mean ± SEM (n = 2 mice). (C) Projection strength to eSCNs vs. CTRNs. Mean ± SEM (eSCNs: n = 4 mice; CTRNs: n = 2 mice).

### Layer-specific targeting of contralateral SC

Commissural SC neurons are thought to play a role in coordinating activity between the two SCs (Takahashi et al., 2005, 2007, 2010). To broaden our understanding of these inter-SC projection patterns, we examined the cell-type-, layer- and, rostrocaudal-specificity of inputs from the contralateral SC. We found that eSCNs received more input from the contralateral SC_id_, as well as the superficial SC, than iSCNs (Wilcoxon rank sum test [one-tailed]; n = 6 mice [eSCNs]; n = 4 mice [iSCNs]; SC_id_: p = 0.0048; superficial SC: p = 0.0095; Fig. 7A-C). We also observed that the rostrocaudal position of contralateral SC_id_ inputs mirrored the rostrocaudal position of starter neurons (Fig.7D-F), such that rostral poles of the two SCs were preferentially interconnected, as were caudal segments. Together, these results suggest that commissural SC neurons mainly target eSCNs located in analogous rostrocaudal positions.

**Figure 7:**
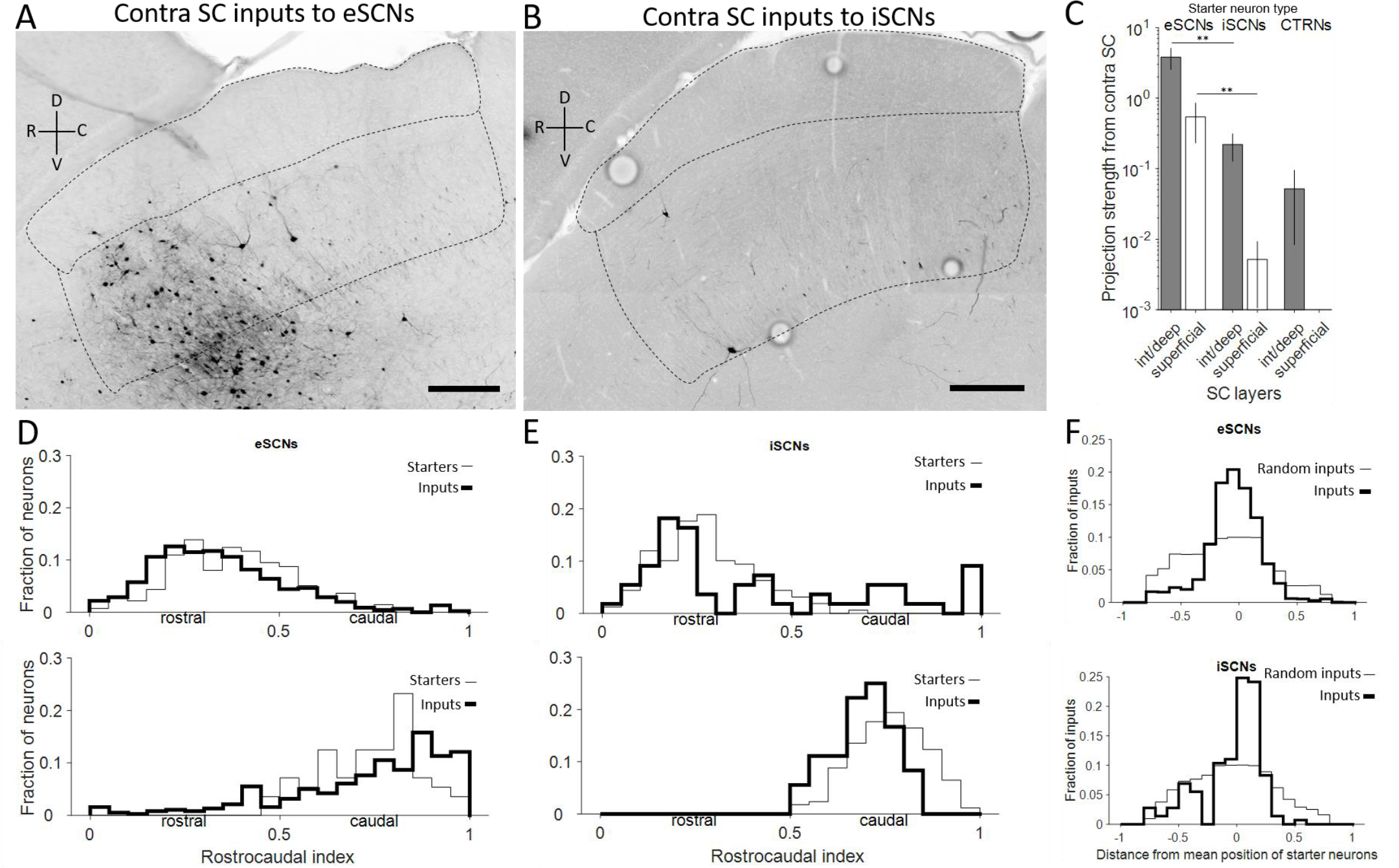
Layer-specific targeting of contralateral SC. (A-B) Contralateral SC inputs to eSCNs (A) or iSCNs (B). Dashed lines depict the borders of the superficial SC, SC_id_, and underlying periaqueductal gray. (C) Projection strength from the contralateral SC_id_ and superficial SC to eSCNs (n = 4 mice), iSCNs (n = 6 mice), and CTRNs (n = 2 mice). Mean ± SEM; **: p<0.01. (D) Normalized rostrocaudal position of excitatory SC_id_ starter and input neurons in the contralateral SC_id_ for two Vglut2-Cre mice. (E) Same as in D except for inhibitory SC_id_ starter neurons in two Gad2-Cre mice. (F) Normalized rostrocaudal distance between input neurons and mean position of starter neurons in each mouse, compared to expected distance for randomly distributed input neurons. eSCNs (top): n = 4 mice; iSCNs (bottom): n = 6 mice.

## 4. DISCUSSION

The SC is critical for a wide range of functions, ranging from spatial attention to multisensory integration to adaptive behavioral responses (Dean et al., 1989; Stein and Stanford, 2008; Gandhi and Katnani, 2011; Krauzlis et al., 2013; Basso and May, 2017). Its intricate internal circuitry is organized into seven cytoarchitechtonically-defined layers composed of excitatory and inhibitory neurons each with diverse projection patterns (Edwards, 1977; Mize, 1992; Olivier et al., 1998, 2000; Pettit et al., 1999; Takahashi et al., 2005, 2007, 2010; Isa and Hall, 2009; Sooksawate et al., 2011; Ghitani et al., 2014). SC_id_ computations are also influenced by the dense innervation from a network of brain structures involved in sensory and motor functions. While this complex anatomical arrangement presents challenges to identify subcircuits for putative SC functions, our study makes inroads into this challenge by employing monosynaptic tracing from identified neuronal subtypes within the SC_id_ and identifies a number of thematic input patterns to the structure varying along a functional axis of rostrocaudal organization. We found that eSCNs received more inputs than iSCNs, extrinsic inputs to eSCNs and iSCNs had a rostral bias, CTRNs were targeted by many fewer brain areas than the general population of eSCNs, and populations of commissurally connected SC neurons were located in similar rostrocaudal positions. While our study has implications for the full range of SC_id_ functions, we focus here on how our findings inform the contributions of the SC_id_ to orienting movements and multisensory integration.

Orienting behaviors can be conceptualized as two discrete components: selecting a target from among multiple options and terminating a target-directed movement appropriately to acquire the target. While these functions are coordinated by computations performed within a network of interconnected brain areas, the dual maps of visual and movement space within the SC_id_ make it a model system in which to study the components of orienting behavior. The superficial layers of the colliculus contain a map of visual space inherited from its direct retinal inputs, raising the question of whether the map of movement space contained within SC_id_ similarly arises from the orderly arrangement of its inputs. For example, does the SC_id_ receive movement commands from the superficial layers or from outside the SC which are then relayed to downstream motor structures, or do critical intrinsic processes within the SC_id_ produce the map of movement space that ultimately determines the vector of the executed movement?

Our findings that different populations of SC neurons receive unique patterns of input argue against the view of the SC_id_ as a simple relay and instead support the idea that intrinsic processing gives rise to SC_id_ computations for orienting behavior. Studies examining SC_id_ activity when multiple targets are present (Basso and Wurtz, 1997; McPeek and Keller, 2004; Li and Basso, 2005; Felsen and Mainen, 2008) support a model of SC_id_ function whereby a “competition” takes place between two or more active foci within the colliculus (Basso and May, 2017). While this competition could simply reflect differences in the activity level of different inputs, local inhibitory connectivity within and between the two SC_id_s (Takahashi et al., 2005, 2007, 2010; Isa and Hall, 2009; Sooksawate et al., 2011; Ghitani et al., 2014) suggests that active intrinsic SC_id_ processes may be at work. Potential roles of inhibition include mediating the competition between multiple regions of SC_id_ representing movement vectors to available targets and sharpening and refining the activity needed to acquire a chosen target. Our finding that iSCNs receive fewer inputs than eSCNs indicates that whichever processes they mediate, a smaller subset of inputs relative to eSCNs is used. Further, our observations that CTRNs receive far fewer inputs than the general population of eSCNs argues against the SC_id_ acting as a relay, and instead suggests that these output neurons are likely sampling and transforming information processed within the SC. Indeed, CTRNs have been found to receive commissural inputs from both eSCNs and iSCNs (Takahashi et al., 2005, 2007, 2010). Thus, our findings support the view that the SC is a critical node in the network of interconnected brain regions responsible for spatial decision making.

Target selection and acquisition are unique components of orienting behavior and are therefore likely to be modulated by distinct inputs. Target selection requires information pertaining to the relative position(s) of one or multiple targets in space, as well as predicted value(s) associated with each target, while acquiring a target will require information with both a high degree of spatiotemporal resolution and up-to-date information on the state of the effector(s) that will be used for acquiring the target. Notably, the strongest projections we observed arise from brain areas well-equipped to provide the various forms of information needed for both target selection and acquisition. Visual, auditory, and barrel cortex are among the regions projecting most strongly to the SC_id_ (Fig. 3) and are likely important for localizing potential targets in space, while the SNr sends a robust inhibitory projection (Graybiel and Ragsdale, 1979), which likely conveys the necessary values associated with individual targets that are required to select among them (Handel and Glimcher, 2000; Basso and Wurtz, 2002; Sato and Hikosaka, 2002; Bryden et al., 2011). Additionally, cerebellar projections to the SC_id_ may be critical in conveying predictive information regarding the position of effectors throughout the trajectory of the movement (Ohyama et al., 2003; Shadmehr, 2017; Owens et al., 2018; Becker and Person, 2019), ultimately mediating successful target acquisition. Indeed, studies performing muscimol inactivation of the cFN in monkeys making saccades to visual targets concluded that cerebellotectal projections might provide the SC_id_ with information about the displacement needed to acquire spatial targets (Goffart et al., 1998). In addition to cerebellar inputs to the SC_id_, our finding that eSCN and iSCN inputs tend to favor the rostral pole of the SC_id_ is also likely to play a role in mediating the small movements required for target acquisition.

The observation that eSCNs receive more inputs than iSCNs suggests that they may also receive a higher degree of convergent inputs, which has implications for how the SC_id_ might process behaviorally relevant multisensory information. Previous work has shown that SC_id_ neurons receive convergent visual, auditory, and somatosensory input (Meredith and Stein, 1986). These multisensory inputs synergistically drive SC_id_ spike output (Miller et al., 2015), which is thought to underlie the saliency of biologically relevant stimuli, allowing animals to produce appropriate orienting responses (Stein and Stanford, 2008; Stein et al., 2014). Modeling work suggests that the supralinear response to multisensory information is attributable to local inhibitory input (Miller et al., 2015). Thus, our observation that iSCNs receive far fewer inputs than eSCNs, and are therefore also less likely to receive convergent multisensory information, allows us to imagine a system whereby fast-spiking iSCNs (Sooksawate et al., 2011) receiving unisensory information contribute to the multisensory enhancement of eSCNs via a local disinhibitory mechanism.

This study has extended our knowledge of SC_id_ inputs and has critically elucidated their cell-type targeting. While technique-specific caveats exist, they do not clearly challenge our interpretations. First, while G-deleted rabies virus-mediated monosynaptic tracing is a powerful tool capable of labeling direct inputs to populations of genetically defined neurons (Wickersham et al., 2007; Callaway and Luo, 2015; Luo et al., 2018), the transmission efficiency can vary based on the molecular composition of presynaptic proteins (Callaway and Luo, 2015). This should be considered when comparing projection strengths across different brain areas. However, since the transmission efficiency is thought to be affected minimally by differences in the cell-type from which the virus is “jumping”, this caveat does not affect our findings that eSCNs receive more inputs than iSCNs and CTRNs, that there is a rostral bias of inputs to eSCNs and iSCNs, or that commissurally connected SC neurons are located in similar rostrocaudal positions.

There are several potential avenues through which future studies can build upon our findings. For example, learning-dependent developmental changes in synaptic connectivity can be examined by combining monosynaptic rabies tracing with a behavioral approach in juvenile mice. Previous work in owls showing that corrupted visual information leads to a topographical misalignment of auditory and visual information within the OT (Brainard and Knudsen, 1998) suggests that task-relevant changes in SC_id_ connectivity might take place during learning. Additionally, similar to the rostrocaudal organization of the SC_id_ map of movement space, there is a mediolateral organization that governs approach vs. avoidance behaviors (Dean et al., 1986, 1989; Sahibzada et al., 1986). While we did not address this in our study, future experiments may wish to examine the extent to which these areas receive unique or overlapping inputs. Finally, the behavioral role of specific inputs to the SC_id_ can be examined by using a self-inactivating rabies virus (Ciabatti et al., 2017) to transport a calcium indicator (Osakada et al., 2011) to reveal input-specific population level neuronal activity during task performance.

**Table 1.** List of anatomical abbreviations used in this study

|  |  |
| --- | --- |
| AI | Agranular insular cortex |
| CbN<sub>I</sub> | Cerebellar nuclei, interposed |
| CbN<sub>L</sub> | Cerebellar nuclei, lateral |
| CbN<sub>M</sub> | Cerebellar nuclei, medial |
| Cin | Cingulate cortex |
| CL | Centrolateral thalamic nucleus |
| CM | Central medial thalamic nucleus |
| DLG | Dorsal lateral geniculate nucleus |
| IGL | Intergeniculate leaflet |
| LO | Lateral orbital cortex |
| LPMR | Lateral posterior thalamic nucleus, mediorostral |
| LPtA | Lateral parietal association cortex |
| M1 | Primary motor cortex |
| M2 | Secondary motor cortex |
| MDL | Mediodorsal thalamic nucleus, lateral |
| MGV | Medial geniculate nucleus ventral |
| MPtA | Medial parietal association cortex |
| OPC | Oval paracentral thalamic nucleus |
| PaF | Parafascicular thalamic nucleus |
| PaXi | Paraxiphoid nucleus of thalamus |
| PC | Paracentral thalamic nucleus |
| PIL | Posterior intralaminar thalamic |
| Po | Posterior thalamic nuclear group |
| PoT | Posterior thalamic nuclear group, triangular |
| PPTg | Pedunculotegmental nucleus |
| PrG | Pregeniculate nucleus of the prethalamus |
| PtPD | Parietal cortex, post, dorsal part |
| PtPR | Parietal cortex, post, dorsal part |
| Reth | Retroethmoid nucleus |
| RRe | Retroreuniens nucleus |
| Rt | Reticular nucleus (prethalamus) |
| S1 | Primary somatosensory cortex |
| S1BF | Primary somatosensory cortex |
| S1FL | Primary somatosensory cortex |
| S1HL | Primary somatosensory cortex |
| S1Sh | Primary somatosensory cortex |
| S1Tr | Primary somatosensory cortex |
| S2 | Secondary somatosensory cortex |
| SNr | Substantia nigra *pars reticulata* |
| SPF | Subparafascicular thalamic nucleus |
| str | Superior thalamic radiation |
| SubG | Subgeniculate nucleus of the prethalamus |
| VM | Ventromedial thalamic nucleus |
| VO | Ventral orbital cortex |
| VPM | Ventral posteriomedial thalamic |
| VPPC | Ventral posterior nucleus parvicellular |
| ZI | Zona incerta |

## ACKNOWLEDGEMENTS

We thank Nathan D. Baker for help with histology and imaging. Light microscopy was performed at the University of Colorado Anschutz Medical Campus Advance Light Microscopy Core supported in part by Rocky Mountain Neurological Disorders Core Grant Number P30NS048154. This work was supported by the NIH/NINDS (R01NS079518) and NIH (R01NS084996).

